# Growth of *Paracoccus aminovorans* is restricted by L-ascorbic acid (LA) and human gut isolate

**DOI:** 10.1101/2025.06.04.657967

**Authors:** Kiran Heer, Pratanu Kayet, Nandita Sharma, Surajit Basak, Saumya Raychaudhuri

## Abstract

Gut microbiota plays a crucial role in the outcome of cholera. In this context, *Paracoccus aminovorans*, a gut commensal, has been evidenced to positively influence the pathogenesis of cholera bacterium. In the present study, we examined the impact of various non-antibiotic strategies on the growth of *Paracoccus aminovorans*. We observed the growth of *Paracoccus aminovarans* was severely restricted in the presence of L-ascorbic acid as well as in co-culture growth assays with another gut commensal, *Weissella confusa*.

## Introduction

*Paracoccus aminovorans* is an obligate aerobe, Gram-negative bacterium that belongs to the genus *Paracoccus*. The genus comprises of 34 species. Members of this genus play a crucial role in the soil microbiome and contribute immensely to the process of microbial remediation of contaminated soil [1, 2]. Intriguingly, *P. aminovorans* is also a member of the human gut microbiota, and it exerts a positive influence on the survival of *Vibrio cholerae*, a causative agent of deadly diarrheal disease cholera, thus contributing to disease progression during outbreak [3, 4]. As *P. aminovorans* aids in the establishment of cholera pathogenesis in the host gut, we wanted to control the growth of this abettor by exploiting some non-antibiotic therapeutic strategies. In this regard, we selected recent therapeutic approaches that are effective against *Vibrio cholerae* under various laboratory conditions. In one such approach where a combination of glucose and probiotic strains effectively restricts *V. cholerae* pathogenesis [5-7], we exploited a similar synbiotic approach with different gut commensals against *P. aminovorans*. In the present study, we isolated commensal bacteria from the fecal sample of a healthy donor and characterized those isolates by molecular taxonomy and genome sequence. One isolate, *Weissella confusa*, in the presence of glucose, appeared to restrict *P. aminovorans* growth in a co-culture growth condition. In the second approach, we exploited L-ascorbic acid (LA), that is effective against *Vibrio cholerae* [8]. We observed that LA also exerts a strong growth inhibitory effect on *P. aminovorans*. Collectively, the present study suggested that non-antibiotic strategies that are effective against *Vibrio cholerae* are equally effective against *P. aminovorans*.

## Materials and methods

### Genome Assembly and Annotation

Raw sequencing data in FASTQ format was obtained via Illumina (Hiseq 2500), 150PE. FastQC and Trimmomatic [9] was used to perform quality control on sequencing reads, effectively removing adaptor sequences, low-quality bases, and short reads to ensure high-quality data for downstream analysis. Excellent-quality trimmed reads were assembled into draft genomes using both SKESA [10] and Spades [11], taking advantage of their excellent efficiency with short-read sequencing data. SKESA excels at producing accurate and contiguous assemblies, whereas Spades provides a robust multi-step assembly strategy. The generated draft genomes underwent additional refinement to ensure a high-quality reconstruction. The quality of the assembled genomes was analyzed using QUAST [12], which offered metrics such as N50, genome length, and misassembly rates, allowing the most reliable assembly to be selected. To determine the closest reference genome, the assembled genome was compared to publically accessible sequences using Mash, a fast genome distance estimator, and BLASTn for sequence similarity searches. The NCBI database provided reference genomes with high sequence identity. The assembled genome was aligned to the reference genome using BWA-MEM v2 [13] with default parameters, and Ivar [14] was used to create consensus. The assembled genome was functionally annotated using Prokka [15], a technique for accurately predicting coding sequences, non-coding RNAs, and other genomic components. The eggNOG-mapper [16] was used to classify proteins based on their function, providing information about gene ontology, metabolic pathways, and evolution. Then we used Barrnap (github.com/tseemann/barrnap) to detect ribosomal RNA (rRNA) genes, whereas Aragon [17] was used to detect transfer RNA (tRNA) and transfer-messenger RNA (tmRNA) genes.

### Antimicrobial Resistance and Horizontal Gene Transfer

To identify antimicrobial resistance (AMR) genes, the genome was analyzed using a combination of AMRFinderPlus [18], RGI (Resistome Gene Identifier) [19], and ResFinder [20]. A comprehensive catalog of resistance factors was obtained. Then we used AlienHunter (www.sanger.ac.uk/tool/alien_hunter/), to identify horizontal gene transfer (HGT) regions. Furthermore, CRISPRCasFinder [21] was used to discover putative CRISPR arrays and Cas genes, suggesting possible interactions between mobile genetic elements and host immunity.

### Mobile Genetic Element Analysis

The assembled genome was then analyzed to find mobile genetic components that contribute to genome plasticity and horizontal gene transfer. ISEScan [22] was used to detect insertion sequences, which indicates the presence of transposable elements and their distribution throughout the genome. ICEberg v3.0 [23] was used to identify integrative and conjugative elements, focusing on regions important in gene transfer and genome evolution.

### Secondary Metabolite

The secondary metabolite biosynthesis gene clusters were analyzed using the Antibiotics and Secondary Metabolite Analysis Shell (antiSMASH v7.0) [24] server with ‘relaxed’ detection strictness. These secondary metabolites play crucial roles in microbial defense, interspecies communication, and have potential therapeutic applications. By using a ‘relaxed’ detection setting, we prioritized the identification of gene clusters with a broader range of predicted biosynthetic pathways.

### Bacterial strains and media

Bacterial strains used for this study are *Paracoccus aminovorans* DSMZ-8537, *Weissella confusa* and *Pediococcus pentosaceus*. Bacterial media used in this study were Luria-Bertani broth (LB) (Bacto, USA), Tryptic soy broth without dextrose (TSB) (Bacto, USA), MacConkey agar (Difco, USA) and de Man, Rogosa and Sharpe agar (MRS) (HiMedia, India).

### Bacterial isolation

For bacterial isolation from fecal samples, freshly voided feces were collected from the volunteers, placed into sterile sealable plastic containers and transported to the laboratory for processing within 2–4 h of collection. Dilutions of fecal samples were made in phosphate-buffered saline (PBS) buffer (pH 7.4), streaked on MRS agar and incubated aerobically for 16-24 h at 37°C.

### Coculture study and spotting assay

*P. aminovorans, W. confusa* and *P. pentosaceus* were grown in TSB medium with different concentrations of glucose up to mid-logarithmic phase. Cocultures (1:1) were set up in TSB containing 0.5%, 1% and 2% glucose. Monocultures were also set up in a similar manner as a control. After 16 h of incubation at 37 °C viability of each culture was checked by making serial dilution in phosphate-buffered saline (PBS) and plating on MRS agar and MacConkey agar. pH of all the cultures was measured at the end point. Plates were photographed after 16 h of growth at 37°C.

### Effect of L-ascorbic acid on growth of *P. aminovorans, W. confusa* and *P. pentosaceus*

To check the effect of L-ascorbic acid on growth of *P. aminovorans, W. confusa* and *P. pentosaceus*, all strains were grown in TSB with 1% glu with increasing concentrations of L-ascorbic acid (0.5 mg/mL, 2.5 mg/mL, and 5 mg/mL, 10 mg/mL and 20 mg/mL) for 16 h at 37°C. Cultures grown were serially diluted and spotted on LB (*P. aminovorans)* and MRS (*W. confusa* and *P. pentosaceus*) agar plates. pH values of each culture were also measured before spotting on agar.

### Effect of cell free culture supernatant (CFCS) on growth of *P. aminovorans*

CFCS of *P. aminovorans, W. confusa* and *P. pentosaceus* was obtained by growing the cultures in TSB with different concentrations of glucose (0.5%, 1.0% and 2.0%) for 16 h at 37°C. After 16 h of incubation cultures were pelleted down, culture supernatant was collected and then passed through 0.22µm syringe filters. Then *P. aminovorans* was grown in its own CFCS (as control) and CFCS of *W. confusa* and *P. pentosaceus* and its survival was checked by spotting of serially diluted cultures on LB agar plates. pH of all the cultures was also checked before spotting on agar.

## Results

### Genomic characterization of *W. confusa* and *P. pentosaceus* isolated from the fecal samples of a healthy donor

*W. confusa* exhibits antimicrobial properties against *V. cholerae* [25]. Literature also suggests probiotic characteristics of *W. confusa* [26]. To isolate this organism and other commensal strains, the fecal sample from a healthy donor was serially diluted on MRS agar plates. Pure colonies were obtained through serial plating, and further taxonomical confirmation of two commensal strains, namely *W. confusa*, and *P. pentosaceus*, were done initially by 16SrDNA sequence analysis. To gain further insight into their genomes, we performed draft genome sequencing. After assembly of the genomes, we got a best N50 score of 198257 and 364106 for *W. confusa* and *P. pentosaceus*, respectively. The final draft genome is approximately 2211942 bp in length with 43% GC for *W. confusa* and 1832229 bp in length with 34.85% of GC for *P. pentosaceus* genome. From genome annotation analysis we observed that *W. confusa* has a total set of 2116 proteins, and detected one tmRNA, 84 tRNA **[Fig:1A]**. Furthermore, we also detected two AMR genes,(e.g. *vanT* gene in the vanG cluster and *vanY* gene in the vanB cluster), one CRISPR array, and 23 HGT sites **[Fig:1B]**. Our analysis of mobile genetic elements detected three insertion sequence (IS) elements and 55 ISE elements in 31 proteins. In the case of *P. pentosaceus* the genome has a total set of 1795 proteins and detected one tmRNA, 55 tRNA **[Fig:1D]**. In addition, there are two AMR genes (*vanT* gene in vanG cluster and *qacG* gene), two CRISPR arrays, and 17 HGT sites **[Fig:1E]**. Our analysis of mobile genetic elements detected 4 insertion sequence (IS) elements and 59 ISE elements in 38 proteins. Our analysis of *W. confusa’s* secondary metabolites found a unique T3PKS (Polyketide Synthase) cluster spanning 41,167 nucleotides and containing 29 genes **[Fig 1C]**. This finding highlights the strain’s ability to produce novel bioactive chemicals. In *P. pentosaceus*, we found three different secondary metabolite biosynthesis clusters: a T3PKS cluster (41,167 nt, 42 genes), a lanthipeptide-class-III cluster (16,196 nt, 12 genes), and a RiPP-like cluster (15,092 nt, 13 genes) **[Fig:1F]**. These clusters indicate a vast genetic arsenal for producing different and potentially novel bioactive compounds, underscoring *P. pentosaceus*’ metabolic adaptability.

**Figure 1:**
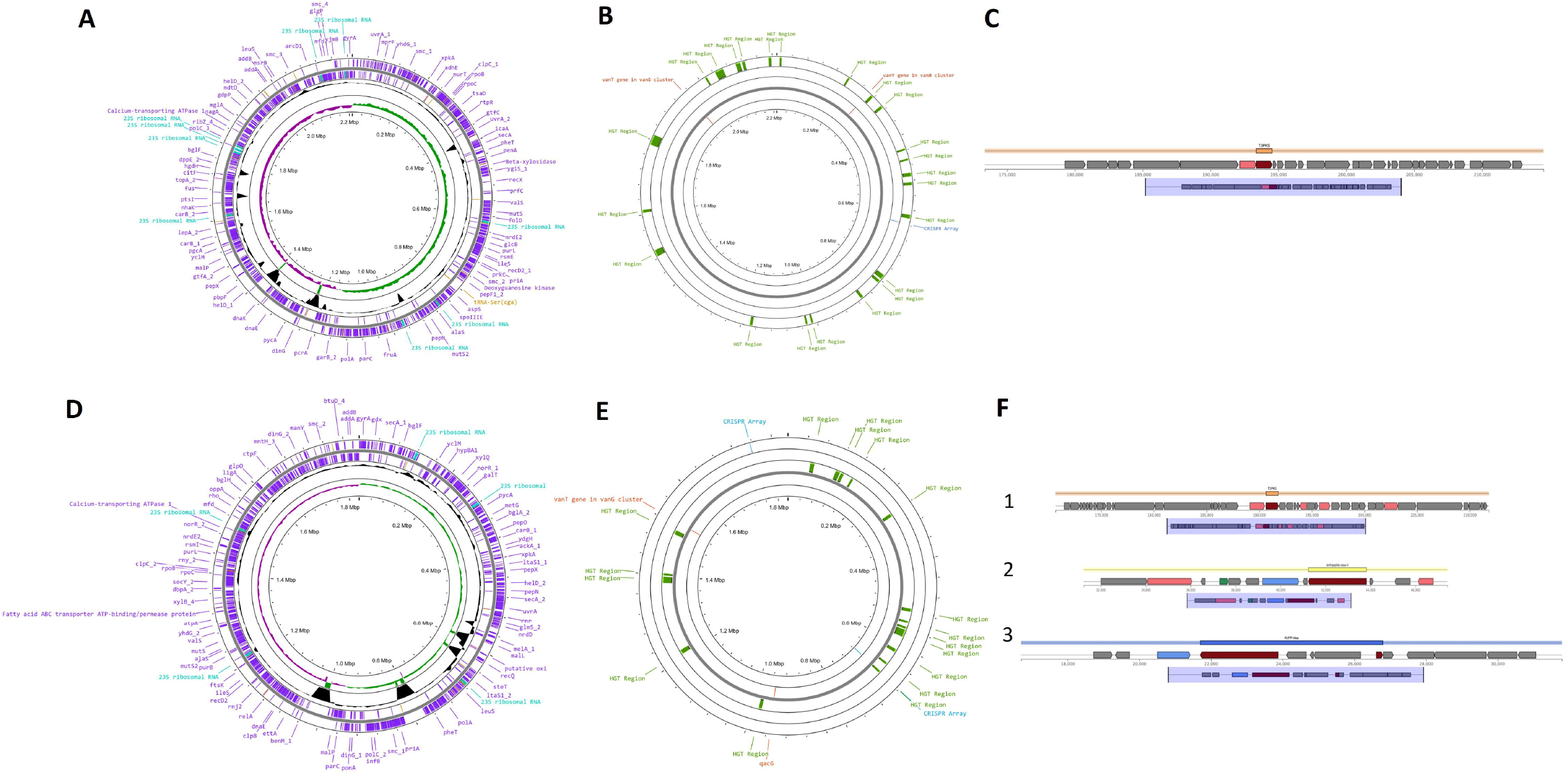
Genomic map of *Weissella confusa* **(A)** where purple colored are genomic CDS and cyan colored are rRNAs and yellow colored are tRNAs. **(B)** red colored are AMR genes and blue colored are CRISPR array and green colored are HGT region. **(C)** Secondary metabolite biosynthesis cluster in *W. confusa*. The T3PKS (Type III Polyketide Synthase) cluster spans 41,167 nucleotides and encodes 29 genes (Genes are depicted in arrow boxes). Genomic map of *Pediococcus pentosaceus* **(D)** where purple colored are genomic CDS and cyan colored are rRNAs and yellow colored are tRNAs. **(E)** red colored are AMR genes and blue colored are CRISPR array and green colored are HGT region. **(F)** Three secondary metabolite biosynthesis clusters in *P. pentosaceus*. (Gens are depicted in arrow boxes) [1] A T3PKS (Type III Polyketide Synthase) cluster spanning 41,167 nucleotides and encoding 42 genes [2] A lanthipeptide-class-III cluster spanning 16,196 nucleotides and encoding 12 genes [C] A RiPP-like (Ribosomally synthesized and Post-translationally modified Peptide) cluster spanning 15,092 nucleotides and encoding 13 genes.

### Weissella confusa mediated growth retardation of Paracoccus aminovorans in the presence of glucose

Both complete 16SrDNA and draft genome analyses confirmed the identity of isolates as *W. confusa* (**Figure 1**) In a similar fashion, we also isolated and characterized another commensal known as *Pediococcus pentosaceus* (**Figure 1**). Previously, *Vibrio cholerae* growth was restricted by glucose and *gut commensals* [5, 6]. This prompted us to examine the effect of the combination of glucose and *W. confusa* as well as glucose and *P. pentosaceus*. Towards this end, *P. aminovorans* was grown in a coculture growth condition either with *W. confusa or P. pentosaceus* in Tryptic soya broth (TSB) with varying glucose concentrations. All the strains exhibited good growth in TSB with glucose **(Figure 2A**). To select each strain in co-culture, we employed different agar plates. For example, McConkey agar selects *P. aminovorans* over *W. confusa* and *P. pentosaceus*, whereas MRS agar promotes the growth of *W. confusa* and *P. pentosaceus* and inhibits the growth of *P. aminovorans* (**Figure 2A**). Our viability spotting assay results indicated growth retardation of *P. aminovorans* by *W. confusa*. No significant growth inhibition is caused by *Pediococcus pentosaceus* in similar growth conditions (**Figure 2B**). In essence, *W. confusa*, which exhibits growth antagonism against *V. cholerae*, is also effective against *P. aminovorans*.

**Figure 2:**
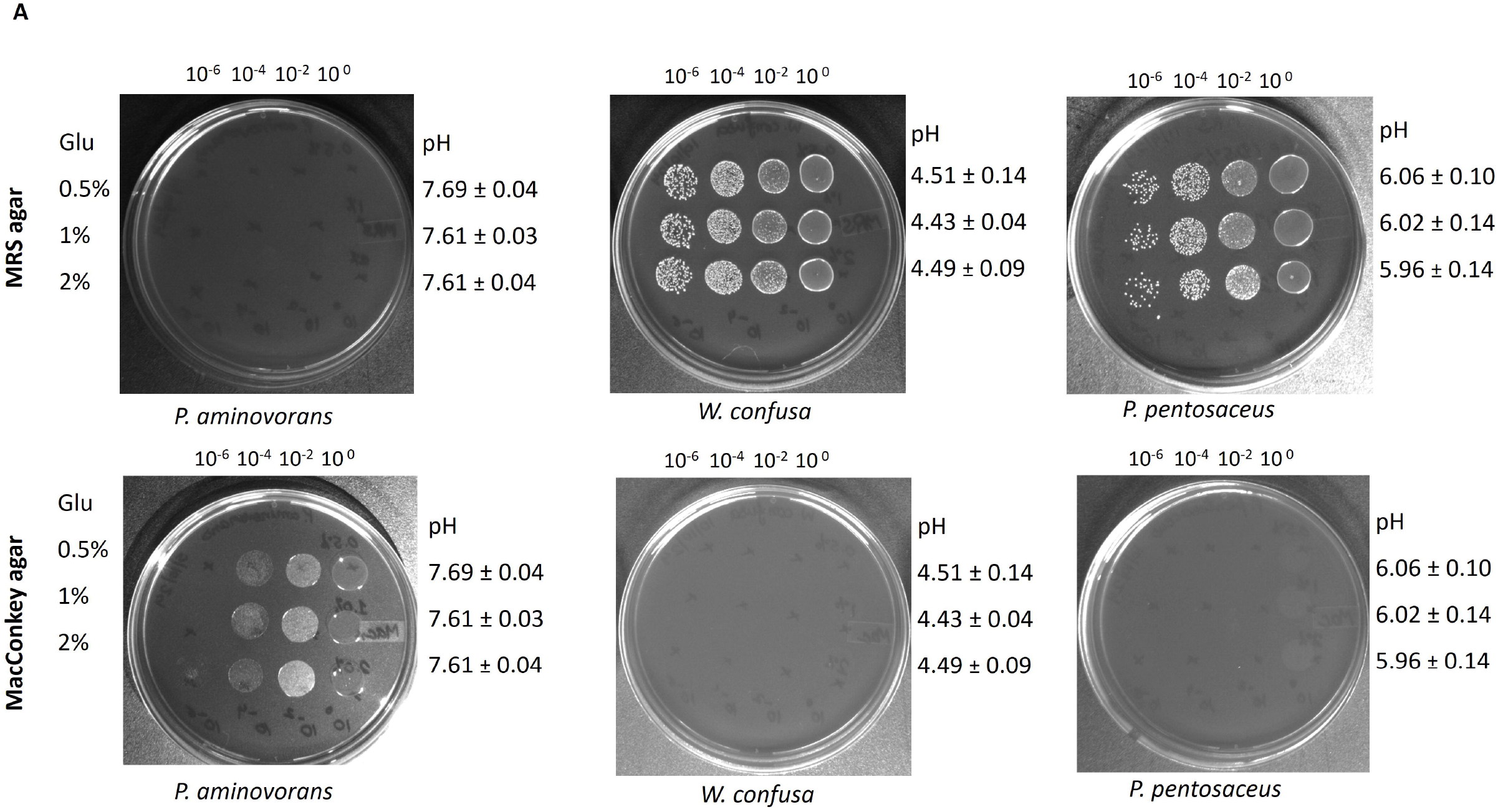
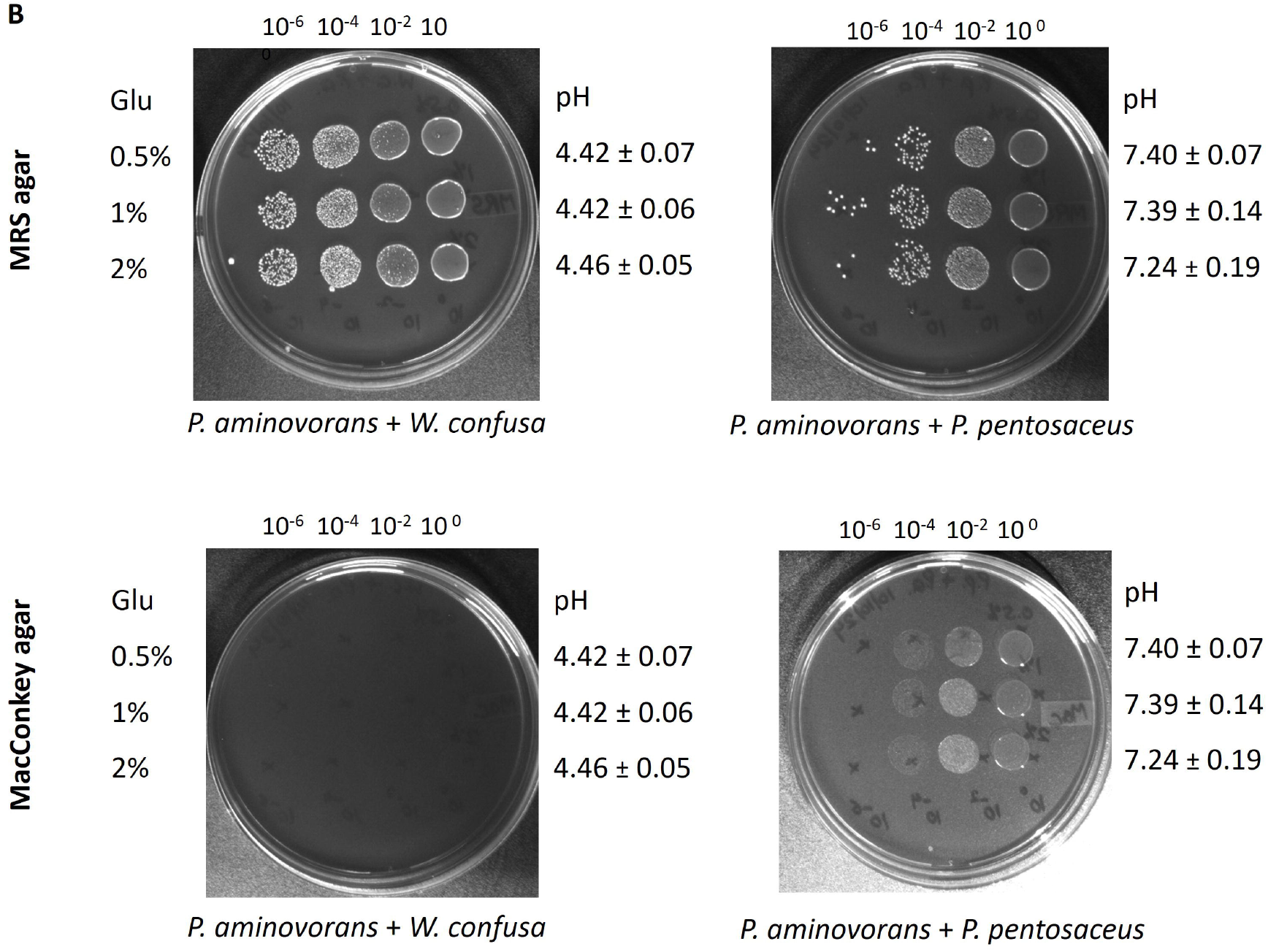
Coculture and spotting assay. **(A)** Spotting of monocultures of *P. aminovorans, W. confusa* and *P. pentosaceus* grown in TSB with different concentration of glucose (0.5%, 1.0% and 2.0%) on MRS agar and MacConkey agar. **(B)** Spotting of coculture of *P. aminovorans* with *W. confusa* and *P. pentosaceus* grown in TSB with different concentration of glucose (0.5%, 1.0% and 2.0%) on MRS agar and MacConkey agar. Cultures grown were serially diluted and spotted on solid agar plates. Plates were photographed after 16 h of incubation at 37 °C. The plate photographs are representative of the experiment performed in biological and technical duplicates. pH values were measured at the end of the growth assay and their standard deviations were rounded off and reported.

### L-ascorbic acid (LA) restricts the growth of Paracoccus aminovorans under in vitro conditions

Recently, L-ascorbic acid (LA) has shown some promise in controlling *Vibrio cholerae* pathogenesis [8]. Keeping in view earlier observations, we wanted to investigate whether LA can control the growth of *P. aminovorans*. Towards this end, *P. aminovorans* was exposed to varying concentrations of L-ascorbic acid in Tryptic soya broth (TSB), which contained 1% glucose. Viability spotting after the stipulated period of incubation evidenced severely reduced growth of *P. aminovorans* with increasing concentrations of LA (**Figure 3**). In the case of *W. confusa*, growth was reduced at a high concentration of LA while *P. pentosaceus* showed an intermediate growth restriction pattern (**Figure 3**). Collectively, we observed that *P. aminovorans* growth can be restricted by gut isolates and LA.

**Figure 3:**
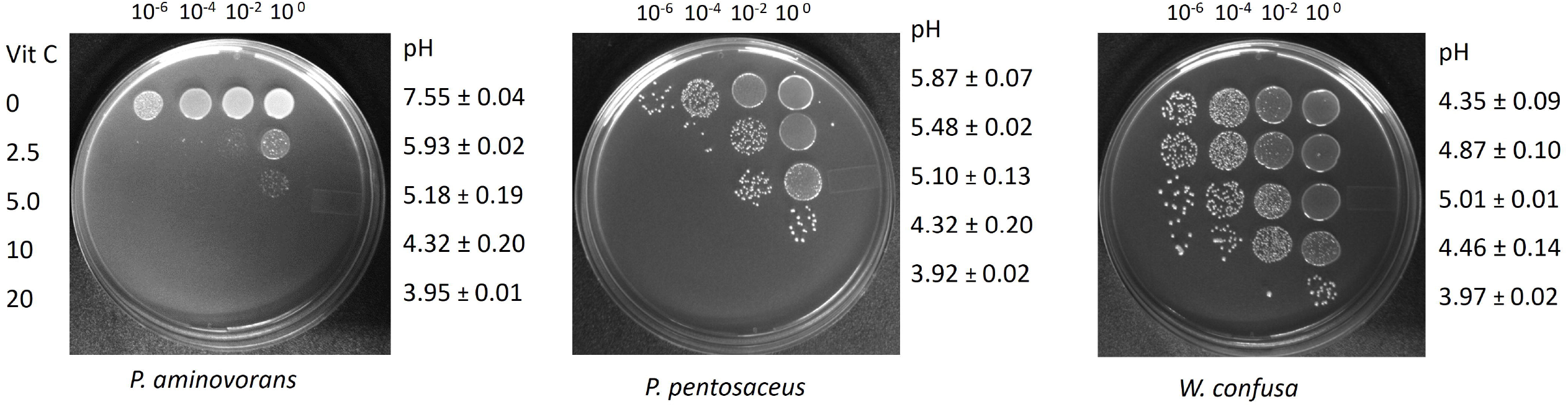
Effect of L-ascorbic acid on growth of *P. aminovorans, W. confusa* and *P. pentosaceus*. All strains were either grown in TSB with 1% glu or TSB with 1% glu supplemented with increasing concentrations of L-ascorbic acid (0.5 mg/mL, 2.5 mg/mL, and 5 mg/mL, 10 mg/mL and 20 mg/mL). Cultures grown were serially diluted and spotted on LB agar plates. pH values were measured at the end of the culture growth and their standard deviations were rounded off and reported.

### Growth inhibition of *P. aminovorans* by cell-free culture supernatant (CFCS) of *W. confusa*

To evaluate whether the killing of *P. aminovorans* by *W*.*confusa* is a contact-independent phenomenon, cell-free supernatant (CFS) of *W. confusa* and *P. pentosaceus* was tested against *P. aminovorans*. We observed that the growth of *P. aminovorans* was completely inhibited only with the CFS of *W. confusa* (**Figure 4**).

**Figure 4:**
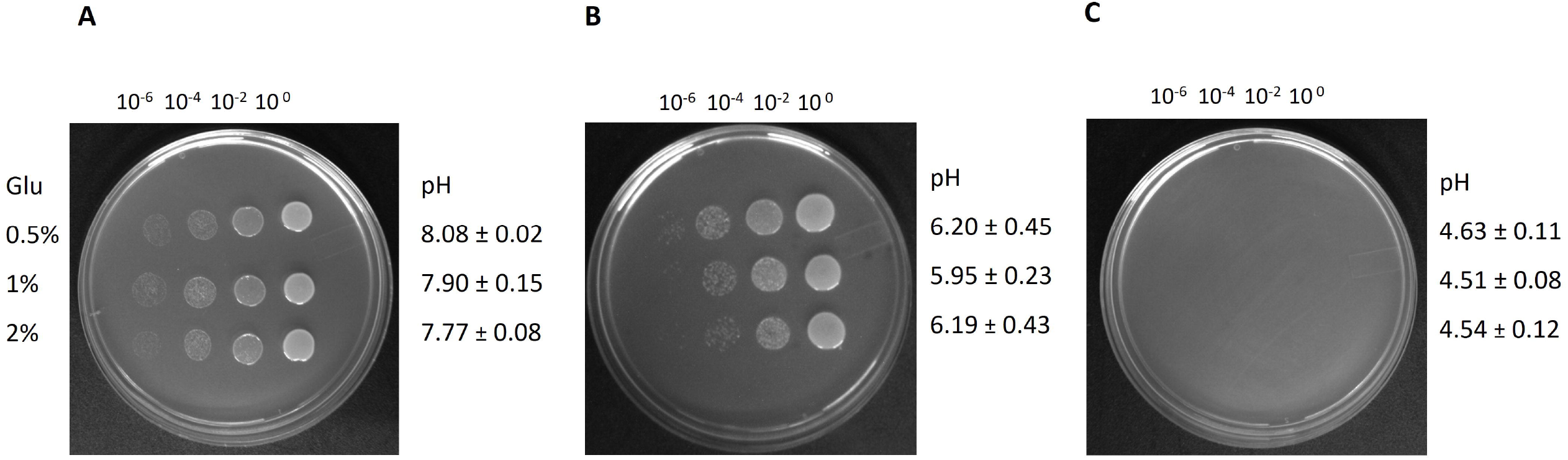
Effect of cell free culture supernatant (CFCS) on growth of *P. aminovorans. P. aminovorans* was grown in CFCS of **(A)** *P. aminovorans*, **(B)** *P. pentosaceus* and **(C)** *W. confusa* and spotting of serially diluted cultures was done on LB agar plates. pH values were measured at the end of the culture growth and their standard deviations were rounded off and reported.

## Discussion

Commensal bacteria play a pivotal role in maintaining and balancing the community structure of the gut environment by providing colonization resistance against pathogens and pathobionts. On the other hand, pathogens and pathobionts are continuously evolving with a panoply of strategies to evade commensal mediated resistance [27, 28]. Interestingly, the role of commensals in aiding pathogenic establishment within the host is also increasingly evident [29]. For example, cross-feeding of sugar-fermented metabolites (e.g. L-lactate) of *Streptococcus gordonii* aids in the pathogenesis of *Aggregatibacter actinomycetemcomitans* [30, 31]. Recently, commensal yeast has been shown to promote the virulence of *Salmonella typhimurium* [32].

The involvement of *P. aminovorans* in the cholera epidemic has paved a new way to understand the disease progression. It is now evident that *Paracoccus* promotes *V. cholerae* colonization in the host intestine [4]. However, there is a paucity of information on the gut dysbiosis caused by *P. aminovorans*. The organism may cause significant dysbiosis through overgrowth (SIBO), and a dysbiotic gut has a direct influence on cholera infection [33, 34]. The possibility of gut dysbiosis caused by *P. aminovorans* should be examined in appropriate animal models that are routinely exploited to study cholera infection [6, 35]. Other than animal models, ex vivo human gut microbiota fecal fermentation models [36, 37] are also suitable for investigating *P. aminovoran*s mediated alteration of community structure. This warrants further investigation.

*V. cholerae* is a remarkably successful pathogen. It has been increasingly recognized how host gut microbiota members modulate the organism’s virulence and survival potential within the host gut ecosystem. For example, *Blautia obeum* (formerly known as *Ruminococcus obeum*) reduces its virulence potential and facilitates host recovery from cholera infection [38]. Similarly, SCFAs produced by gut commensals are also effective in restricting the growth of the organism [39]. On the other hand, *P. aminovorans* facilitates multispecies biofilm development and promotes colonization of *V. cholerae* in the host intestine [4, 40], but the exact mechanism (e.g cross feeding of metabolites) is yet to be explored. Like *P. aminovorans*, other gut community members may also facilitate cholera. It is, therefore, necessary to understand the diarrheal disease in the context of the gut ecology.

## Conclusion

Our present endeavor, suggests the possibility of exploiting *W. confusa* to control the growth of *P. aminovorans*. Recently, the growth of *Vibrio cholerae* has also been found to be inhibited by *W. confusa* [25]. Therefore, the effect of *W. confusa* to control cholera infection should be evaluated in the presence of both *V. cholerae* and *P. aminovorans* in a suitable animal model. Additional studies are necessary to address this issue.

## Declarations

## Abbreviations

*P. aminovorans*: *Paracoccus aminovorans*
*W. confusa*: *Weissella confusa*
*P. pentosaceus*: *Pediococcus pentosaceus*
*V. cholerae*: *Vibrio cholerae*
LA: L-ascorbic acid

## Ethics approval and consent to participate

Approval for study was obtained from the Institutional Ethical Review Committee of CSIR-Institute of Microbial Technology (IEC-IMTECH) (IEC-January 2020#1) and all the guidelines and regulations were followed while performing the experiments. Written consent was obtained from volunteers prior to the collection of fecal matter.

## Consent for publication

Not applicable

## Data availability

The Whole Genome sequences have been deposited at GenBank. The NCBI GenBank accession numbers for *Pediococcus pentosaceus* is JBKIBA000000000 and for *Weissella confusa* JBKIBC000000000.

## Competing interest

All authors declare no conflict of interest.

## Funding

This work was partly supported by grants from Science and Engineering Research Board (CRG/2018/000297/SERB-GAP/0185) and by in-house grants (OLP-151 and OLP-193) and CSIR-MLP 39. KH and NS and acknowledge CSIR for fellowship. The funder has no role in the design of the study, analysis, and interpretation of data and in writing the manuscript.

## Author contributions

SR conceived the idea. SR, and KH planned the experiments. SR wrote the manuscript’s original draft. KH isolated gut commensals and characterized by employing molecular tools. KH performed all experiments. NS repeated spotting assays. PK and SB carried out genomic analysis and annotation. SR, KH and SB edited the final manuscript. All authors gave editorial inputs.

## Acknowledgements

We acknowledge all fecal sample donors. We extend our appreciation to all donors for their support.

## Clinical trial number

Not applicable.

## Notes

### Competing Interest Statement

The authors have declared no competing interest.

